# rRNA depletion for holobiont metatranscriptome profiling across demosponges

**DOI:** 10.1101/2022.08.12.503726

**Authors:** Sergio Vargas, Ramón E. Rivera-Vicéns, Michael Eitel, Laura Leiva, Gabrielle Büttner, Gert Wörheide

## Abstract

Despite the extensive knowledge of sponge microbiome diversity, a critical knowledge gap persists concerning the molecular mechanisms that govern host-symbiont interactions. Deciphering these mechanisms is crucial for understanding how sponge holobionts respond to environmental changes and identifying potential disruptions in sponge-microbe associations. A hindrance to progress in characterizing these molecular crosstalk mechanisms is the scarcity of broadly applicable molecular methods for efficiently sequencing meta-transcriptomes across a diverse array of sponge species. To tackle this challenge, we have introduced a hybrid-capture strategy capable of selectively depleting sponge and bacterial rRNA from total RNA extracts obtained from highly divergent demosponges with varying microbiome complexities. Our innovative pan-demosponge rRNA depletion approach streamlines the efficient characterization of metatranscriptomes within diverse demosponge holobionts, concurrently facilitating the quantification of gene expression in both the host and its microbiome. This methodological advancement represents a significant stride in unraveling the molecular intricacies of sponge-microbe interactions, providing a robust platform for future investigations across a broad spectrum of sponge species.

## Introduction

With almost 8,000 species, class Demospongiae is the most diverse sponge class comprising ~70% of all described species. Demosponges occur at all depths in all oceans, and representatives of this sponge class can also inhabit lakes and rivers of all continents except Antarctica [1].

One of the most striking aspects of the biology of demosponges is their pervasive association with bacterial communities of varying diversity and composition [2]. First discovered and described in the late 1970s, the study of sponge-bacteria associations has evolved into a new and vastly expanding field. More than 40 bacterial phyla have been associated with sponges, forming the sponge holobiont [3]. Importantly, these bacterial associates differ from the bacterioplankton of the surrounding water and sediments and include sponge-specific bacterial clades [3, 4]. Despite the large body of information available on sponge microbiome diversity, the molecular mechanisms used by sponges to interact with their bacterial symbionts and *vice versa* remain largely unexplored [5, 6]. This knowledge is needed, for example, to appreciate better how changes in the symbiotic communities in response to short-term or long-term environmental change affect the sponge host.

The lack of experimentally tractable sponge models has made the study of sponge-bacteria associations difficult [7]. Besides, the lack of broadly applicable methods for functional genomics of demosponges has limited the study of sponge holobionts to a few species. Thus, generalizations about the biology of demosponge holobionts are based on the few species for which genomic or (meta-)transcriptomic resources are available [8–12]. Consequently, developing molecular methods applicable to a broad taxonomic spectrum of demosponges to, for instance, simultaneously characterize the transcriptome of the sponge host and its microbiome - under control and experimental setups - is pivotal. Sponge nucleic acid extracts frequently are complex mixtures of DNA/RNA derived from the sponge and the community of sponge-associated micro- and macro-organisms. Although careful sample preparation can minimize the risk of coextraction for most macro-organisms, separating the sponge cells from associated microorganisms is usually difficult without compromising sample quality. Thus, sponge total DNA/RNA extracts typically contain molecules derived from the host and its microbial associates.

Studies involving transcriptomics typically use two messenger RNA (mRNA) enrichment strategies before library preparation and sequencing: polyA-capture and rRNA depletion (Fig. 1). Enriching for mRNA before library preparation and sequencing is necessary due to the stark difference in abundance between mRNAs, transcriptomics’ target molecules, and rRNAs, the most abundant RNAs in total RNA extractions. In yeast, for instance, one microgram (1 μg) of total RNA contains only around 50ng of mRNA, with most remaining RNA being rRNA and tRNA [13, 14]. Thus, directly sequencing total RNA will mainly yield rRNA and tRNA sequences, with abundant mRNA only appearing in low counts at prohibitive sequencing depths and rare mRNAs absent altogether. PolyA-capture leverages poly-dT oligonucleotides to selectively transcribe or capture mRNAs before proceeding with library preparation and sequencing. The method is widely used for eukaryotic material as it is easy to implement and typically results in mRNA-enriched samples suitable for transcriptomics. However, since it relies on the presence of a polyA tail in the target mRNA molecules, it is unsuitable for bacterial transcriptomics. rRNA-depletion does not require polyadenylated mRNAs, but knowledge about the sequence of the rRNA molecules to be depleted is necessary, limiting its application to previously characterized samples and making it challenging to deplete rRNA from complex mixtures such as sponge-derived total RNA extracts.

**Figure 1.**
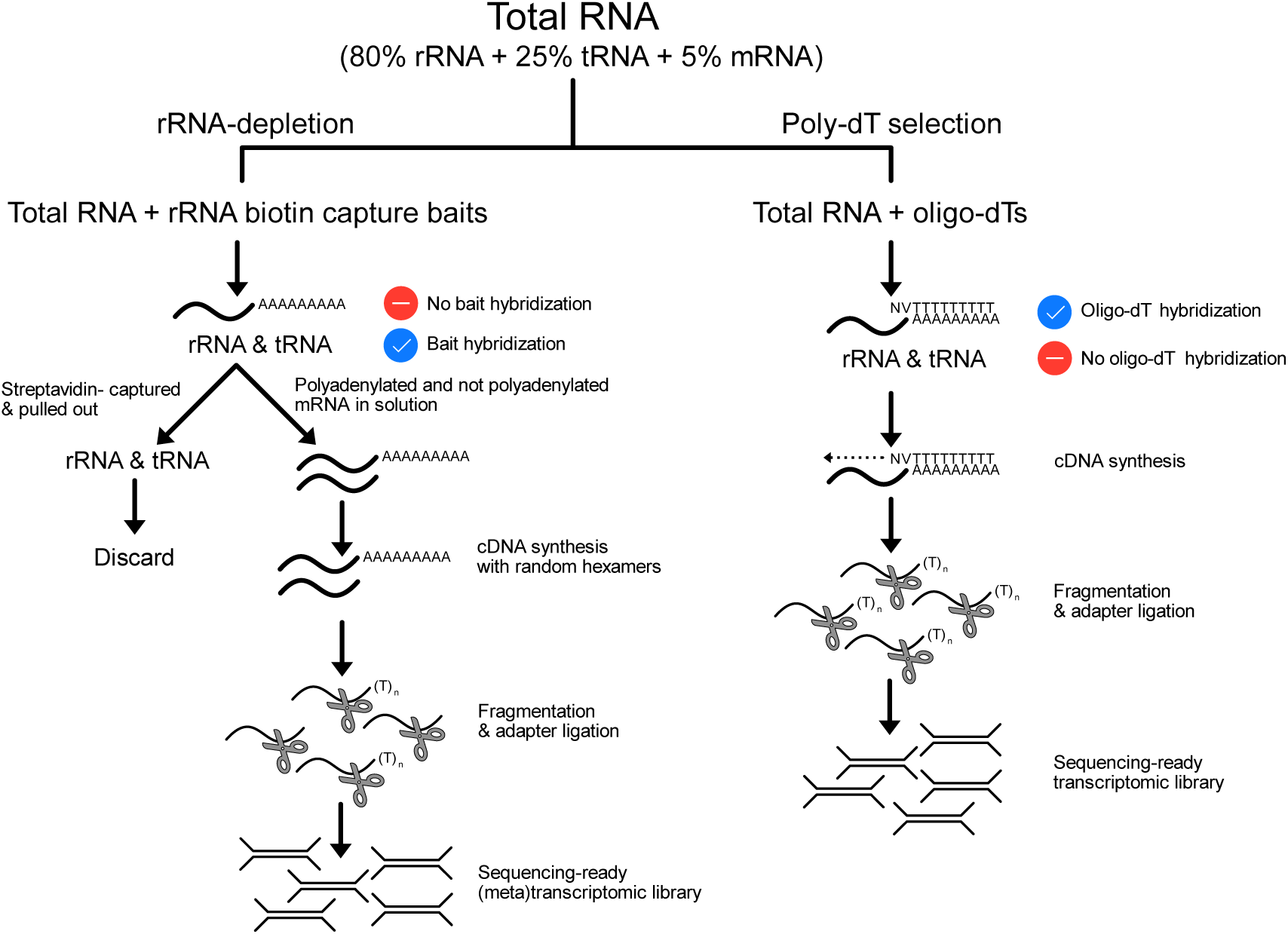
Messenger RNA (mRNA) enrichment strategies and sample preparation for transcriptome sequencing. In polyA selection (right), oligo-dTs serve either as capture baits or as primers for retrotranscription, thereby producing mRNA-enriched solutions; rRNAs and tRNAs are not primed/captured by oligo-dTs and are discarded during the procedure. Contrary to polyA-selection, rRNA depletion (left) captures rRNAs with complementary baits and discards the captured molecules, leaving polyadenylated and non-polyadenylated mRNAs in solution for downstream analyses. Depletion-based strategies enable metatranscriptome sequencing because (polyadenylated) eukaryotic and (non-polyadenylated) prokaryotic mRNA remains available for downstream processing.

Here, we present a pan-demosponge rRNA capturing strategy capable of depleting the rRNAs of phylogenetically diverse sponge hosts and their associated bacterial community. As a proof of concept, we use this procedure to sequence the meta-transcriptomes of *Lendenfeldia chondrodes* and *Tethya wilhelma*, two demosponge species with contrasting microbiomes: *Lendenfeldia chondrodes* is a high microbial abundance keratose sponge with a microbiome dominated by cyanobacteria while *T. wilhelma* is a low microbial abundance sponge belonging the demosponge subclass Heteroscleromorpha. Keratosa and Heteroscleromorpha diverged >700MYA [15] and represent the phylogenetic diversity breadth of Demospongiae. Finally, we evaluate whether rRNA-depleted libraries better represent the expressed bacterial gene set and can be leveraged to study differential gene expression of the sponge microbiome.

## Materials and Methods

### Selection of 28S and 18S rRNA demosponge sequences for pan-demosponge depletion bait design

Full-length nuclear 18S, 5.8S, and 28S and mitochondrial 12S and 16S rRNA sequences from the demosponges *Amphimedon queenslandica, Tethya wilhelma, Lendenfeldia chondrodes,* and *Spongilla lacustris* (Accession numbers OX176765-OX176794) were processed by siTOOLs Biotech (Germany) using a proprietary bait selection technology to produce a cocktail of 106 pan-demosponge ribosomal depletion baits. The resulting pan-demosponge bait set can be purchased directly from siTOOLs Biotech (Germany). Supplementary Table 1 details which sequences were used for each marker by species.

### Bleaching experiments in *Lendenfeldia chondrodes*

Round explants (1cm in diameter) were shaded in the same 360L marine aquarium (25 °C water temperature and a 12:12 h light cycle) where the donating sponges were cultured. We used black plastic foil mounted on plexiglass panels to cover the aquarium space and shade the explants. Neither the plastic foil nor the plexiglass panels were in contact with or blocked the water (in-/out-)flow. Control and treatment (*i.e.*, shaded) explants were placed on (open) Petri dishes (14cm diameter; Carl Roth AG, Germany), separated to avoid contact (and fusion), and visually inspected every week to check for signs of necrosis or negative effects of the treatment. Petri dishes were cleaned weekly from accumulating sand or fine sediment using a Pasteur pipette. This process took less than one hour and did not interfere with the shading treatment. Due to the symmetrical arrangement of the water inflow in the experimental tank and the presence of a single, central outflow, both control and treatment sponges were exposed to similar temperatures, nutrients, water pH, and water currents, thus allowing to constrain these potential confounding effects from the experimental setting. Whole explants were fixed in liquid nitrogen after 12-15 weeks of exposure to darkness, avoiding transferring other organisms found in the aquarium, like small amphipods or polychaetes. Two 1cm diameter control (unshaded) explants were fixed in liquid nitrogen the same day the shading treatment ended. Tissue samples were kept at −80 °C until DNA/RNA extraction.

### Total RNA extraction in Lendenfeldia chondrodes and Tethya wilhelma

RNA from flash-frozen treatment (n=2) and control (n=2) *L. chondrodes* (see above) and one flash-frozen control *T. wilhelma* specimen was extracted using Trizol. Briefly, the frozen tissue was submerged in Trizol and homogenized using a Polytron PT1300D homogenizer. We vigorously mixed the homogenate for five minutes at room temperature and added 200 μl chloroform per ml homogenate for phase separation. We centrifuged the samples at maximum speed (21,000 RCF) for 15 minutes and transferred the upper phase to a new microfuge tube for precipitation with isopropanol. After centrifugation (21,000 RCF at 16 °C), we washed the pellet twice with ethanol 80%, air-dried it, and resuspended it in nuclease-free water. We assessed RNA yield and purity using a Nanodrop 100 and controlled the quality of the extracts using agarose gels (1%) before checking RNA integrity on a Bioanalyzer 2100.

### qPCR-based abundance estimation of Cyanobacteria and Actinobacteria in control vs. bleached *Lendenfeldia chondrodes*

We used a qPCR-based relative quantification to determine the abundance of Cyanobacteria and Actinobacteria in control *vs.* bleached *L. chondrodes* sponges. We selected Actinobacteria because specific primers previously tested in sponges are readily available [16], and we expected this non-photosynthetic taxon to be insensitive to the shading treatment in contrast to the photosynthetic Cyanobacterium. Furthermore, OTUs belonging to these bacterial phyla occur in *L. chondrodes* [17]. We extracted DNA from control (n=3) and bleached (n=3) *L. chondrodes* explants using the NucleoSpin® Tissue kit (Macherey-Nagel). We used 15uL qPCR reactions of the Kappa SYBR Fast Ready-Mix (Kappa Biosystems) to conduct, for each tissue sample, three qPCR reactions using (1) universal bacterial primers, (2) Cyanobacteria specific primers and (3) Actinobacteria specific primers [16]. We used two technical replicates for all qPCR reactions, no-template negative controls, and a Rotor-Gene Q (Qiagen) RT-qPCR machine for cycling. We analyzed the raw fluorescence measurements obtained for the samples using the program LinReg (https://medischebiologie.nl/) to get Cq and efficiency values for each reaction. We used REST12 [18] to estimate bacterial abundance fold changes and their associated p-values using the mean Cq and efficiency values derived from each reaction’s two technical replicates.

### Ribosomal RNA depletion in Lendenfeldia chondrodes and Tethya wilhelma

Ribosomal RNA depletion from total RNA samples of *L. chondrodes* and *T. wilhelma* was done by siTOOLs using the riboPOOL Buffer Set (Hybridization Buffer: 10 mM Tris-HCl pH 7.5, 1 mM EDTA, 2 M NaCl; Depletion Buffer: 10 mM Tris-HCl pH 7.5, 1 mM EDTA, 1 M NaCl) and magnetic beads. One µg total RNA was incubated with 100 pmol of each pan-bacteria and pan-demosponge (see Supplementary Table 2 for details) riboPOOL probe mixtures roughly determined from the bacterial to sponge RNA ratio observed in total RNA bioanalyzer runs of each of the studied sponge species (Fig. 1). To prepare the riboPOOL mixture, pan-demosponge, and pan-bacteria riboPOOLs were diluted in nuclease-free water to a final concentration of 100pmol/µl (100 µM) and mixed to the desired ratios to a final volume of 10 µl (e.g., 5µl of pan-demosponge riboPOOL with 5µl pan-bacteria riboPOOL for the 50:50 % mix). The hybridization was done by siTOOLs for each sample independently in 0.2 μl thin-wall PCR tubes, using one µg total RNA diluted in 14 μl ddH2O, 5µl Hybridisation Buffer, and one μl 100pmol/µl riboPOOL mix. A non-depletion (“no riboPOOL”) control was performed for samples with enough RNA by replacing the riboPOOL mix with dd H_2_O. The hybridization reaction was incubated at 68 °C for 10 minutes and slowly cooled to 37 °C in a PCR machine with a ramping rate of 3 °C/min. The hybridized rRNA was captured using 540 μl siTOOLs riboDepletion beads washed once with 540 μl Depletion Buffer and resuspended in 480 μl Depletion Buffer. Eighty μl beads were added to each hybridization reaction, and the mix was incubated for 15 minutes at 37 °C followed by five minutes at 50 °C. The capturing reaction was placed on a magnetic rack, and the (clear) supernatant containing the depleted RNA was transferred to a new 1.5 mL tube. The rRNA-depleted samples were purified using the Macherei-Nagel RNA XS Clean-up Kit following the manufacturer’s instructions and eluted in 10µl ddH_2_O. The concentrated, rRNA-depleted samples were quantified in a Biotek EPOCH Photometer with a Take3 Micro-Volume Plate, and the depletion efficiency was controlled on a Bioanalyzer 2100 using the Agilent RNA 6000 Pico Kit (Supplementary Table 3). The depleted RNA was transported on dry ice for library preparation at the Department of Earth and Environmental Sciences, Paleontology & Geobiology, Ludwig-Maximilians-Universität München.

### Library preparation, sequencing, and bioinformatic analyses of rRNA-depleted *Lendenfeldia chondrodes* and *Tethya wilhelma* samples

Lexogen’s CORALL kit was used to construct total RNA (non-polyA-selected) Illumina-ready libraries using one nanogram of the depleted RNA samples, *i.e.*, two control and two bleached *L. chondrodes* samples and one *T. wilhelma*. After library preparation, we quality-controlled and estimated the length of the fragments in the libraries on a Bioanalyzer 2100 using Agilent’s High Sensitivity DNA kit. We sequenced one equimolar pool of the four *L. chondrodes* libraries and the *Tethya* library on an Illumina MiniSeq using one High Throughput 150 Cycles Kit for each species. We *de novo* assembled and annotated the resulting data using TransPi [19], and we used *blast* and *sortmerna* to quantify the number of transcripts matching the Uniprot (trembl) Bacteria database and the amount of host and symbiont rRNA contamination in the sequenced libraries, respectively. To compare the effect of rRNA depletion on the representation of bacterial transcripts in *T. wilhelma* and *L. chondrodes*, we used TransPi to assemble and annotate a similar number of polyA-selected reads obtained from publicly available RNA-Seq datasets (*T. wilhelma*; SRR4255675) or previously sequenced polyA-selected libraries derived from bleached (n=2) and control (n=2) *L. chondrodes* (PRJEB24503). Within TransPi, we evaluated transcriptome completeness using BUSCO v4 for Metazoa and Bacteria [20]. To assess the ability of rRNA-depleted and polyA-selected libraries to detect transcripts from the bacterial symbionts, we used Salmon [21] to map the metatranscriptomic and polyA-selected reads to the predicted protein encoding genes (pegs) of one cyanobacterium and one actinobacterium (Supplementary File 1 and 2, respectively) present in the microbiome of *L. chondrodes* [17]. We then used the resulting count matrices in DESeq2 [22] to assess whether bacterial transcripts are overrepresented in depleted compared to polyA-selected libraries of bleached and control *L. chondrodes* specimens.

## Results

### Pan-demosponge rRNA depletion enriches bacterial transcripts in two phylogenetically distant demosponges with contrasting microbiomes

In both *L. chondrodes* and *T. wilhelma*, rRNA depletion using the newly designed pan-demosponge baits resulted in rRNA-free RNA extracts according to bioanalyzer profiles (Fig. 1). Analyses with *sortmerna* revealed a broad spectrum of rRNA contamination in the depleted libraries with the RNA extracts derived from control *L. chondrodes* sample exhibiting the highest amount of rRNA in the libraries (19% and 27%), followed by the bleached *L. chondrodes* (11% and 19%) samples and *T. wilhelma* (6%); for reference, the pooled polyA-selected *L. chondrodes* library had only 2% rRNA contamination. The assembled meta-transcriptomes for these two species contained 111,891 (with 31,797 >500bp) and 100,018 (33,141 >500bp) transcripts, respectively. For comparison, the polyA-selected transcriptomes for *L. chondrodes* and *T. wilhelma* contained 36,370 (16,754 >500bp) and 53,880 (20,509 >500bp) transcripts, respectively. We also observed an increase in the total (bacterial + metazoan) number of contigs containing ORFs recovered from each species when using rRNA depletion. In this regard, the *L. chondrodes* and *T. wilhelma* transcriptomes assembled from rRNA-depleted RNA had 2.34 and 1.65 times more contigs containing ORFs than their polyA-selected counterparts. The observed increase in transcripts (and contigs containing ORFs) was accompanied by a decrease in the transcriptome N50 and mean transcript length of the metatranscriptome of both species compared to their polyA counterparts (see Table 1). A BUSCO analysis revealed marked differences in the percentage of complete BUSCOs between the rRNA-depleted and polyA-selected transcriptomes in *L. chondrodes* and *T. wilhelma*. Here, the rRNA-depleted *L. chondrodes* and *T. wilhelma* metatranscriptomic assemblies had 73.6% and 70.8% complete metazoan BUSCOs, respectively, while their polyA-selected counterparts had 91.5% and 91.3% complete BUSCOs. This drop in the percentage of complete BUSCOs recovered in the rRNA-depleted assemblies roughly matches the increase in the fragmented BUSCOs observed in these datasets; fragmented BUSCOs accounted for 14.8% and 13.2% of the BUSCOs present in the metatranscriptomes of *T. wilhelma* and *L. chondrodes*, respectively. Regarding bacterial transcriptome completeness, rRNA depletion had a different effect in the two species investigated in this study. In *L. chondrodes*, compared with the polyA-selected assembly, rRNA-depletion resulted in a considerable increase in the complete bacterial BUSCOs recovered. In this species, the number of complete bacterial BUSCOs found was 53.3%, contrasting with the 9.7% bacterial BUSCOs detected in the polyA-selected assemblies. This trend was also evident if the analysis was restricted to Cyanobacteria and Actinobacteria BUSCOs (Supplementary Table 4). However, in *T. wilhelma*, rRNA depletion did not result in a marked increase in the bacterial BUSCOs detected. In this species, we found 19.3% complete bacterial BUSCOs in the rRNA-depleted assembly and 9.7% complete bacterial BUSCOs in the polyA-selected *T. wilhelma* transcriptome. Table 1 summarizes the rRNA-depleted and polyA-selected (non-depleted) assemblies of *L. chondrodes* and *T. wilhelma*.

**Table 1:** Description of *L. chondrodes* and *T. wilhelma* rRNA depleted and polyA-selected transcriptomes. Transcript N50 and mean length in base pairs. *Busco v5 yielded identical results.

|  | *L. chondrodes* |  | *T. wilhelma* |  |
| --- | --- | --- | --- | --- |
|  | rRNA depleted | polyA selection | rRNA depleted | polyA selection |
| N transcripts | 111,891 | 36,370 | 100,018 | 53,880 |
| N50 | 932 | 2,044 | 1,155 | 2,101 |
| Mean length | 555 | 994 | 641 | 901 |
| Total N ORFs (host+ symbiont) | 49,185 | 21,019 | 43,261 | 26,212 |
| Complete/Fragmented BUSCOs (Metazoa v4) | 73.6%/13.2% | 91.5%/1.4% | 70.8%/14.8% | 91.3%/1.8% |
| Complete/Fragmented BUSCOs (Bacteria v4)* | 53.3%/8.1% | 9.7%/4.0% | 19.3%/7.3% | 9.7%/2.4% |

In line with the BUSCO results, blast searches against bacterial sequences deposited in the UniProt knowledge base (*i.e.*, SwissProt + Trembl) revealed a marked increase in the number of bacterial transcripts detected in rRNA-depleted *vs.* polyA-selected transcriptomes in *L. chondrodes* (Fig. 2). In *T. wilhelma*, the difference in the number of transcripts assigned to bacteria in the rRNA-depleted *vs.* polyA-selected transcriptome was less pronounced (Fig. 2).

**Figure 2.**
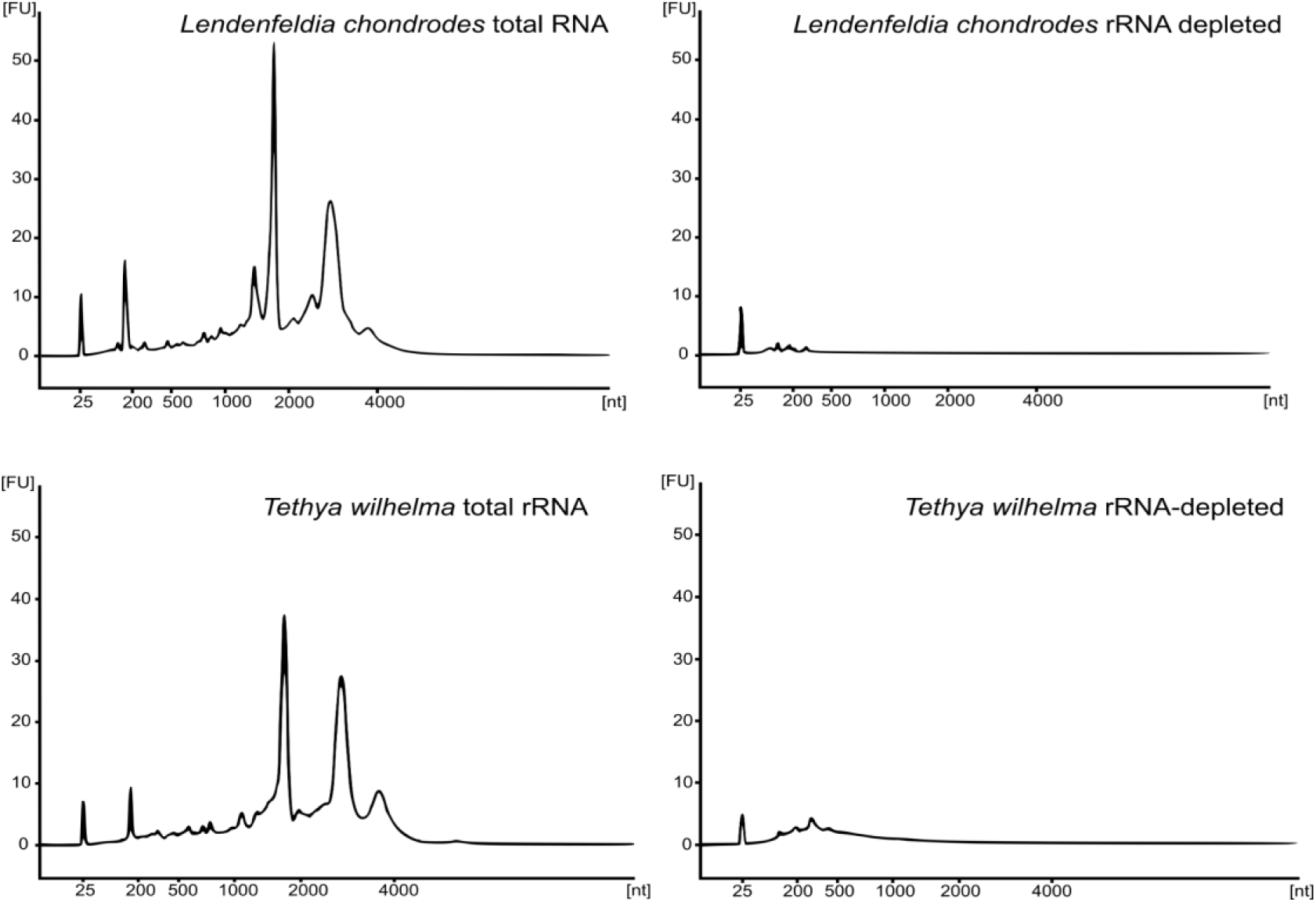
Bioanalyzer electropherograms of “no riboPOOL” input total RNA extracts and riboPOOL depleted control and bleached samples of *Lendenfeldia chondrodes* and samples of *Tethya wilhelma* showing effective rRNA depletion.

### rRNA-depletion enables differential gene expression analyses of the *L. chondrodes* holobiont

rRNA-depleted *L. chondrodes* libraries yielded better bacterial transcript coverage and higher median transcript counts than polyA-selected equivalents. In rRNA-depleted libraries, 5,633 of 5,857 (>99%) bacterial transcripts had at least one read count in at least one of the four libraries sequenced, and 1,820 of 5,857 (32%) bacterial transcripts were detected in all sequenced libraries. In rRNA-depleted libraries, the median count ranged between zero (in one of the bleached specimens) and 25 counts. In contrast, only 1,828 of 5,857 bacterial transcripts (32%) had non-zero counts in at least one of the four polyA-selected libraries analyzed, and only seven transcripts (0.001%) were detected in all four polyA-selected libraries analyzed. The median count for polyA-selected libraries was zero, regardless of whether the libraries derived from bleached or control *L. chondrodes* samples. In agreement with these results, rRNA-depleted libraries yielded broad count distributions that contrasted with those obtained via polyA-selection, which had prominent peaks at low counts, indicating a general depletion of bacterial reads in these libraries (Fig. 3). The depletion of bacterial transcripts (*i.e.*, count peak displaced towards the left in Fig. 3) in the sequenced transcriptomes was evident in bleached *L. chondrodes* independent of the method used for library preparation. However, rRNA-depleted libraries obtained from bleached *L. chondrodes* recovered considerably more (4,918 *vs.* 167) bacterial transcripts than polyA-selected libraries from similar samples.

**Figure 3:**
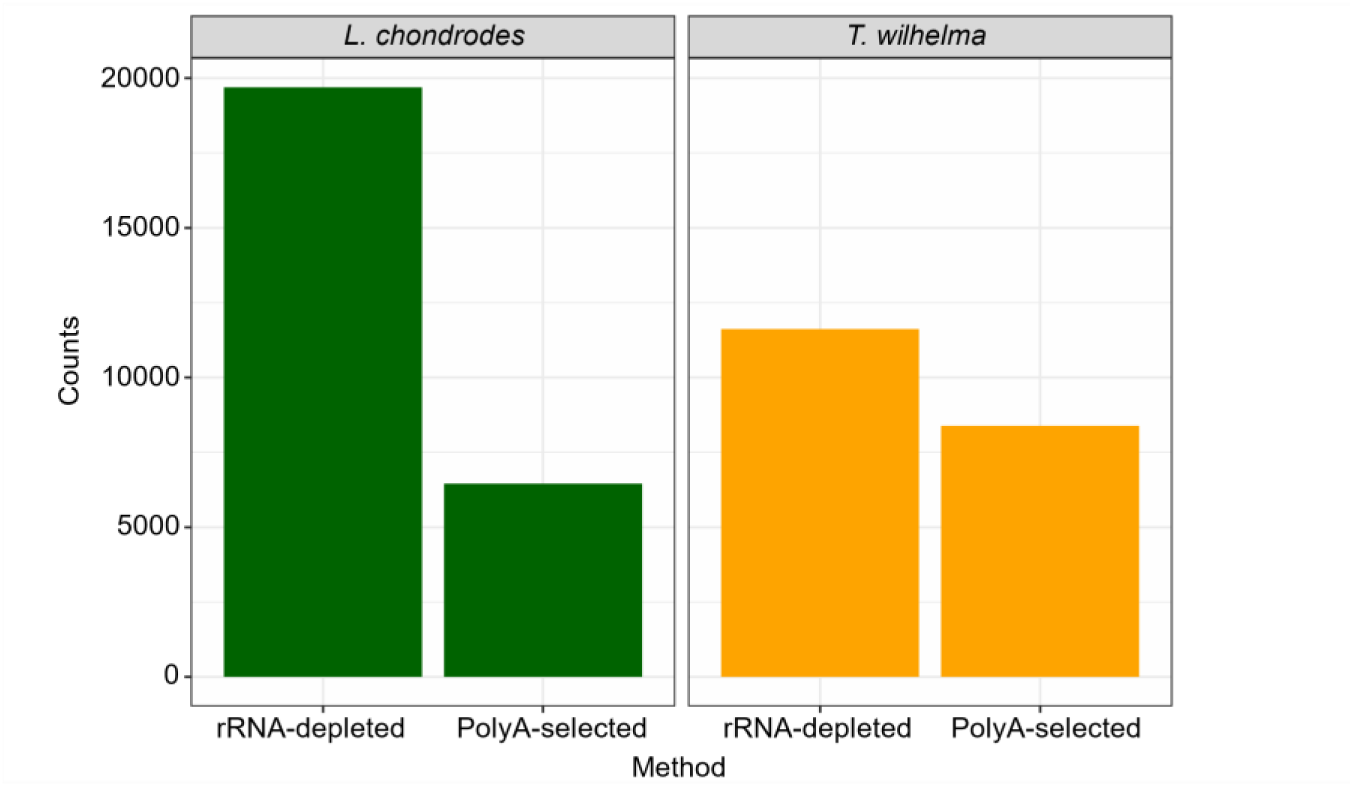
Number of transcripts in the assembled reference transcriptome matching bacterial sequences deposited in the UniProt knowledge base in rRNA-depleted vs. polyA-selected transcriptomes of the demosponges *Lendenfeldia chondrodes* and *Tethya wilhelma*.

Relative quantification assays performed via RT-qPCR (Table 2) revealed a significant negative six log2 fold change in the abundance of cyanobacteria in shaded relative to control sponges. In contrast, the abundance of symbiotic Actinobacteria remained unaffected by shading (Table 2). rRNA-depleted libraries partially mirrored this microbiome change, yielding cyanobacterial transcript count distributions biased towards lower counts in bleached *L. chondrodes* specimens (Fig. 3). In contrast, rRNA-depleted libraries derived from bleached specimens had higher actinobacterial transcript counts (Fig. 3). PolyA-selected libraries generally failed to detect actinobacterial transcripts (Fig. 3).

**Table 2:** REST analysis of the relative abundance quantification of the cyanobacterium Candidatus *Synechococcus spongiarum* and a sponge-associated Actinobacteria in bleached *vs.* control sponge explants. We used 1000 iterations for the calculation of p-values.

| Bacterial Group | Used as | Mean Reaction Efficiency* | Expression | Std. Error | 95% C.I. | P(H1) | Result |
| --- | --- | --- | --- | --- | --- | --- | --- |
| Eubacteria | Reference | 0.963 | 1.000 |  |  |  |  |
| Cyanobacteria | Target | 0.8461 | 0.022 | 0.011 - 0.046 | 0.007 - 0.061 | 0.007 | DOWN |
| Actinobacteria | Target | 0.9614 | 0.589 | 0.254 - 1.339 | 0.165 - 2.351 | 0.300 |  |

\* For the REST calculations, the efficiency was calculated for each reaction using the program LINREG

DESeq2 analyses revealed a generalized increase in the actinobacterial and cyanobacterial counts in depleted libraries. For instance, 1,924 of 2,533 (76%) actinobacterial and 2,029 of 3,100 (65%) cyanobacterial transcripts had at least two times more (log2-fold change ≥ 1, adjusted p-value < 0.05) counts in depleted libraries derived from bleached specimens (Fig. 4). Similarly, in libraries derived from control specimens 1,473 of 3,100 (47%) cyanobacterial transcripts had higher counts (log2-fold change ≥ 1, adjusted p-value < 0.05) in depleted *vs.* polyA-selected libraries. In contrast, only 147 of 2,533 (6%) actinobacterial transcripts had higher counts in depleted *vs.* polyA-selected libraries (Fig. 4). However, this apparent difference in the enrichment of cyanobacterial and actinobacterial transcripts in libraries derived from control specimens was caused by the inability to compute p-values for most (1,476 of 2,533; 58%) actinobacterial transcripts and the high p-values for most remaining transcripts (900 of 1057 transcripts with adjusted p-value ≥ 0.05).

**Figure 4:**
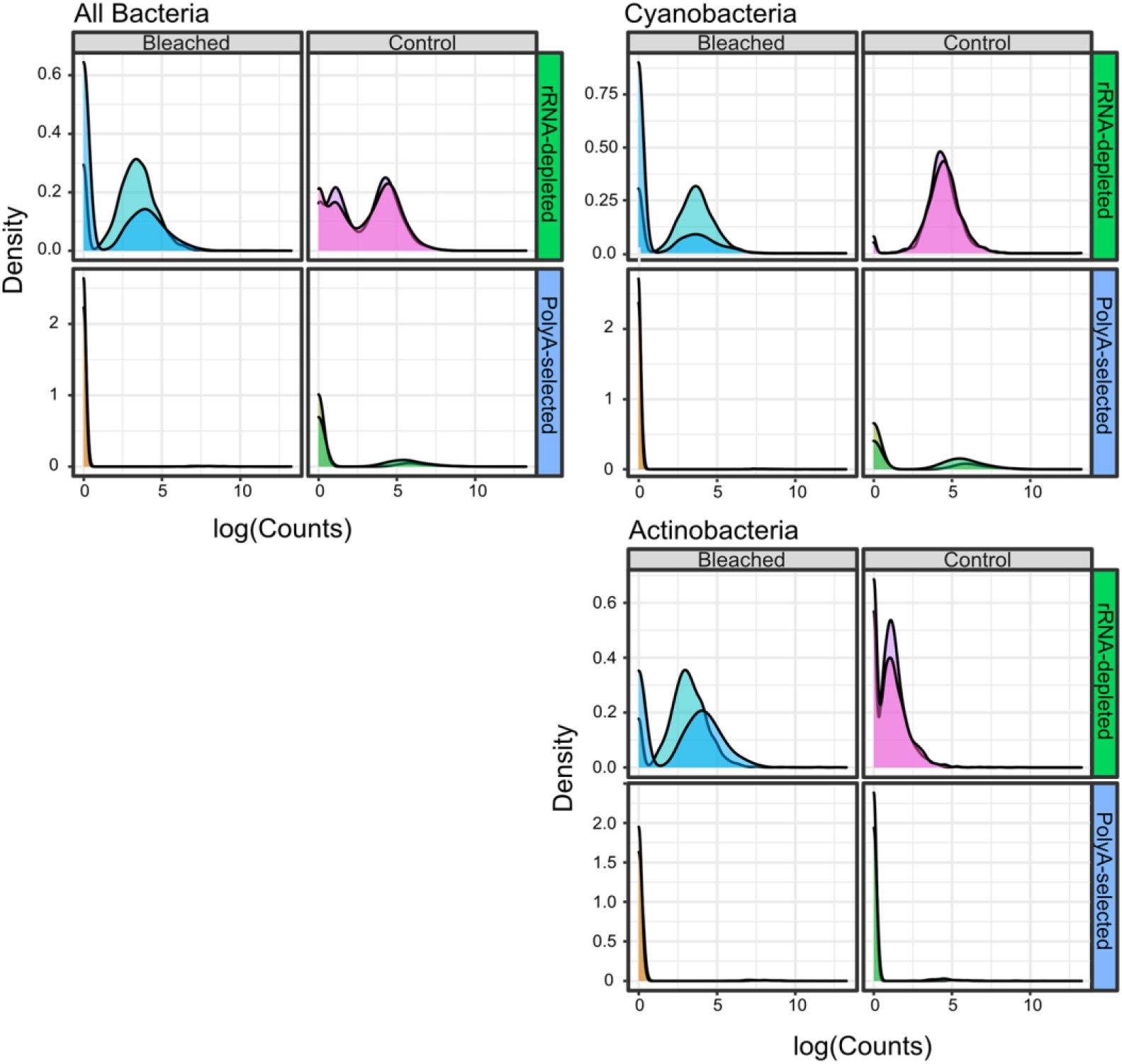
Distributions of log-transformed read counts obtained after mapping rRNA-depleted and polyA-selected control and bleached *Lendenfeldia chondrodes* libraries against bacterial protein encoding genes from two bacteria present in this species. Two control and two bleached sponges were used for polyA-selection and rRNA-depletion, as indicate by the shading levels of the same color.

**Figure 5:**
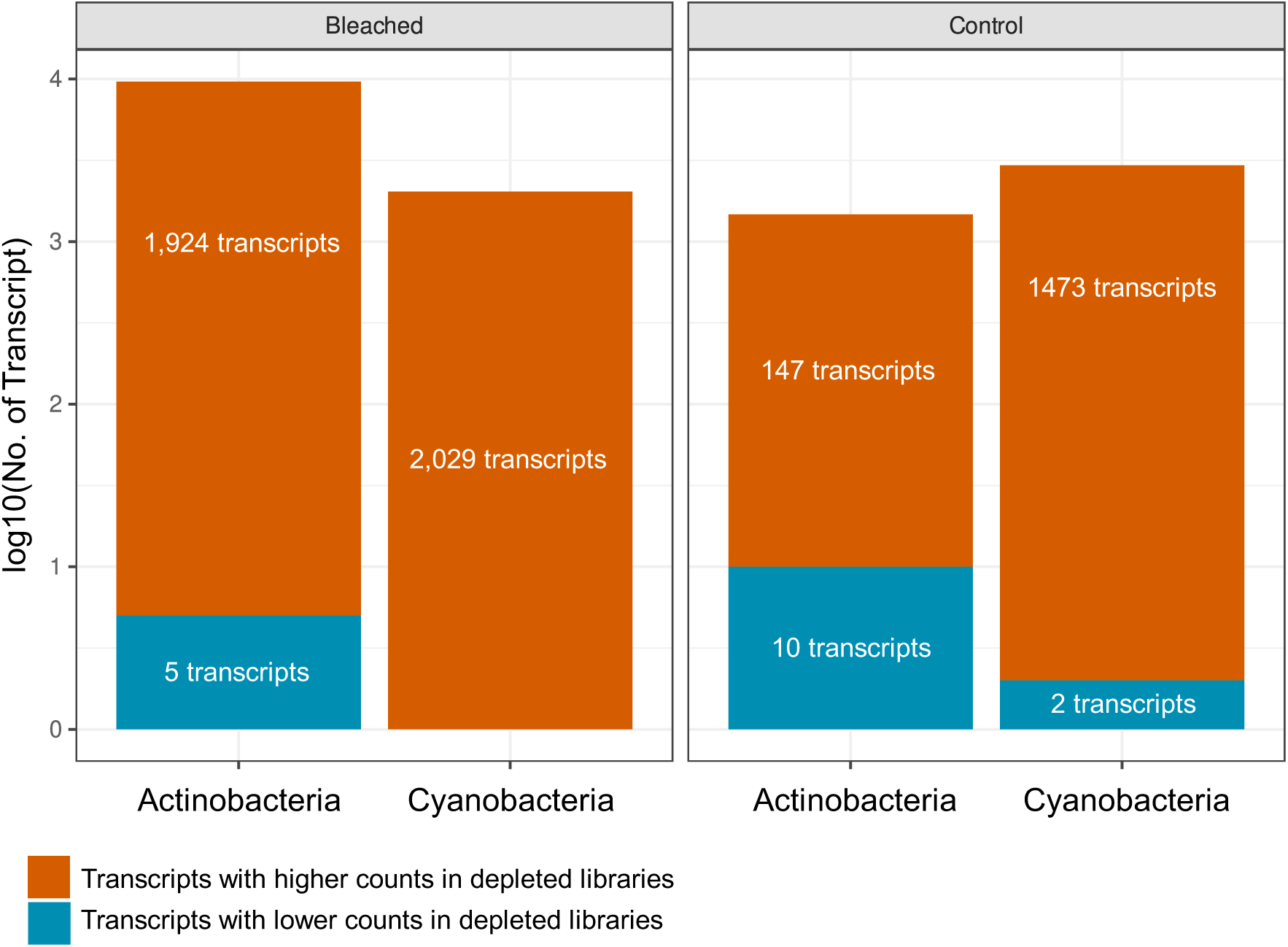
Number of cyanobacterial and actinobacterial transcripts with higher/lower counts in depleted libraries derived from bleached and control *Lendenfeldia chondrodes* samples. To determine whether a transcript had a higher/lower count in depleted libraries the transcript counts observed in polyA-selected libraries was used as the reference level in a DESeq2 analysis.

## Discussion

Sponges and their microbial communities are integrated systems with complex physiologies [5] that, in some contexts, are better regarded as holobionts with physiological and metabolism different from those of their individual constituents. Thus, analytical methods to study how sponges respond to changes due to intrinsic (*e.g.*, ontogenetic changes) and extrinsic factors (*e.g.*, competition, predation by other organisms, or varying environmental conditions) must account for this complexity. Here, we designed a pan-demosponge rRNA-depletion bait set for metatranscriptome sequencing in Demospongiae, the most species-rich sponge class, and tested it in two distantly related demosponge species, namely *T. wilhelma* and *L. chondrodes*. Our rRNA-depletion strategy detected two to five times more bacterial BUSCOs and recovered two times more bacterial protein-coding transcripts than polyA-selected assemblies of these species. Thus, as intended and expected, our library preparation strategy better captures the complex nature of sponge holobionts than polyA-based methods for producing RNASeq libraries.

A reduction in assembly N50 and contig mean length accompanied the increase in protein-coding transcripts, bacterial transcripts, and bacterial BUSCOs. This is expected given the shorter length of prokaryotic compared with eukaryotic genes [23]. Hence, the drop in N50 and contig mean length observed in our metatranscriptomes likely reflects the effect of a larger bacterial gene pool detected in these assemblies on these two metrics, especially considering that depleted libraries were assembled using longer (150bp) reads than polyA-selected libraries. In addition, due to their higher complexity, it is also likely that the rRNA-depleted metatranscriptomic assemblies are more fragmented than their polyA-selected counterparts. In this regard, the increased number of fragmented BUSCOs in the metatranscriptomic assemblies of both *T. wilhelma* and *L. chondrodes* provides evidence for their fragmentation. Considering that rRNA-depleted RNA extracts include non-polyadenylated sponge RNA species (*e.g.*, lncRNAs) and substantially more bacterial mRNA transcripts absent in polyA-selected equivalents, the observed fragmentation of rRNA-depleted compared to polyA-selected assemblies at similar sequencing depths is not entirely unexpected. Using similar read lengths, a relation between transcriptome complexity, assembly quality, and sequencing depth exists [24, 25], and comparisons between multiple non-model animals suggested that ca. 30×10^6^ pairs of reads could consistently generate high-quality metazoan assemblies [25]. We followed this recommendation but obtained metatranscriptomic assemblies that substantially differed from their polyA-selected equivalents, despite using longer reads for depleted metatranscriptomes.

Sequencing depth recommendations available to date are based on the analysis of polyA-selected libraries. Our results, using two sponge species with contrasting microbiomes [17, 26], suggest that bacterial diversity (i.e., the number of species present in a microbiome and their abundances) might influence the quality of the obtained metatranscriptomes. In this regard, *T. wilhelma*’s metatranscriptomic assembly barely differed from its polyA-selected counterpart in the number of complete bacterial BUSCOs, while *L. chondrodes*’ assembly using rRNA depletion showed a substantial increase in the obtained number of complete bacterial BUSCOs compared to the polyA-selected assembly. *Tethya wilhelma*’s microbiome is less complex than that of *L. chondrodes* [17, 26], and the bacteria in *T. wilhelma* occur in lower abundances those in *L. chondrodes*, a species with abundant cyanobacterial symbionts. Assuming a baseline of 30×10^6^ read pairs for polyA-selected metazoan transcriptomes, increasing the number of reads *n* times with *n* equal to the number of holobiont members seems adequate when sequencing rDNA-depleted total RNA. However, the optimal sequencing depth for rRNA-depleted libraries remains to be empirically determined and individually adjusted based on the complexity of the studied holobiont. Cyanobacteria dominate the microbiome of *L. chondrodes* [17], and exposing this species to shading causes the collapse of the cyanobacterial population [27, 28]. As our RT-qPCR relative quantification results indicate, Cyanobacteria were significantly less abundant in shaded *vs.* control *L. chondrodes* sponge samples. On the contrary, Actinobacteria, a non-photosynthetic bacterial phylum in *L. chondrodes*’ core microbiome [17], did not change in abundance upon shading. Compared with polyA-selected libraries, rRNA-depleted libraries derived from control and bleached *L. chondrodes* samples were enriched in actinobacterial and cyanobacterial transcripts. rRNA-depleted libraries derived from control samples were apparently less enriched in actinobacterial transcripts than libraries derived from bleached specimens.

It is likely that the slight enrichment of actinobacterial transcripts in rRNA-depleted control libraries compared with polyA-selected control libraries resulted from the impossibility to calculate p-values for most transcripts caused by the systematic low counts obtained for Actinobacteria in polyA-selected libraries in control samples. In this regard, control samples contain an overwhelmingly high cyanobacterial population that possibly hampers the adequate sampling of transcripts derived from other bacteria. Supporting this idea, we observed higher actinobacterial counts in rRNA-depleted libraries derived from bleached specimens. Given that sequencing depth was similar between rRNA-depleted libraries and that the Actinobacteria species does not change its abundance upon shading, the lower counts obtained for actinobacterial transcripts in control samples likely results from the higher abundance of Cyanobacteria (and their transcripts) in these samples. The better detectability of actinobacterial transcripts due to the decay of the cyanobacterial population indicates that the sequencing depth for metatranscriptomic experiments needs to be determined empirically to compensate for the abundance of different bacterial groups. This is especially important for differential gene expression studies since the interplay between bacterial abundance and transcript detectability can result in artifactual differential gene expression caused by changes in bacterial abundance or composition in the studied sponges. Here, we have presented an rRNA-depletion strategy for simultaneously sequencing the transcriptomes of demosponges and their microbial communities. We tested this strategy in two non-closely related demosponges with contrasting microbiomes and showed its capacity to recover more bacterial transcripts and capture changes in the bacterial transcriptome caused by environmental perturbations (*i.e.*, shading). We consider our strategy an opportunity to standardize the generation of sponge metatranscriptomes, facilitating the metatranscriptomic profiling of sponge holobionts along their ontogeny, the study of their responses to environmental change, and between-species comparisons of these systems. We furthermore emphasize that - following individual taxon-specific adaptations - this approach can be expanded to cover other sponge classes and animal groups.

## Acknowledgments

We thank Andrew Walsh and Dr. Michaela Beitzinger from siTOOLs Biotech for designing the biotinylated DNA oligos and conducting the rRNA depletion experiments, respectively. Michael Hannus and Anna Liznar (siTOOLs) provided important input during the early phases of the project and were open to experimenting with not necessarily high-profit sponges. RERV, ME and GW acknowledge funding from the European Union’s Horizon 2020 research and innovation programme under the Marie Skłodowska-Curie grant agreement No. 764840 (ITN IGNITE). SV thanks N. Villalobos Trigueros, M. Vargas Villalobos, S. Vargas Villalobos, and S. Vargas Villalobos for their constant support. N. Villalobos helped fetching and formatting the bacterial uniprot database using custom Python scripts.

## Data Availability

The metatranscriptomic reads generated for this study are available under Bioproject PRJEB53671 PolyA-selected reads were taken from Bioproject PRJEB24503 and PRJNA288690. The metatranscriptomic assemblies generated with the corresponding annotations, the bacterial metagenomic assemblies used to map metatranscriptomic reads for differential gene expression analyses, and the scripts used are available in the project repository: https://gitlab.lrz.de/cbas/cbas_resources.

## Author contributions

**SV.** Conceptualization, Methodology, Investigation, Resources, Visualization, Writing - original draft. **RERV.** Software, Writing - review and editing. **ME.** Project administration, Writing - review and editing. **GB, LL.** Investigation. **GW.** Resources, Supervision, Funding acquisition, Writing - review and editing.

---

**Supplementary Table 1:** Used rRNA sequences by sponge species included in bait design.

| Species | Nuclear rRNAs |  |  | Mitochondrial rRNAs |  |
| --- | --- | --- | --- | --- | --- |
|  | 18S | 5.8S | 28S | 12S | 16S |
| *Amphimedon queenslandica* | X | - | X | - | X |
| *Ephydatia fluviatilis* | X | X | X | X | X |
| *Spongilla lacustris* | X | X | X | X | X |
| *Lendenfeldia chondrodes* | - | X | X | - | - |
| *Tethya* spp.* | X | X | X | - | - |

\* *Tethya wilhelma* OR *Tethya minuta* sequences were used depending on availability, quality and sequence length.

**Supplementary Table 2:**
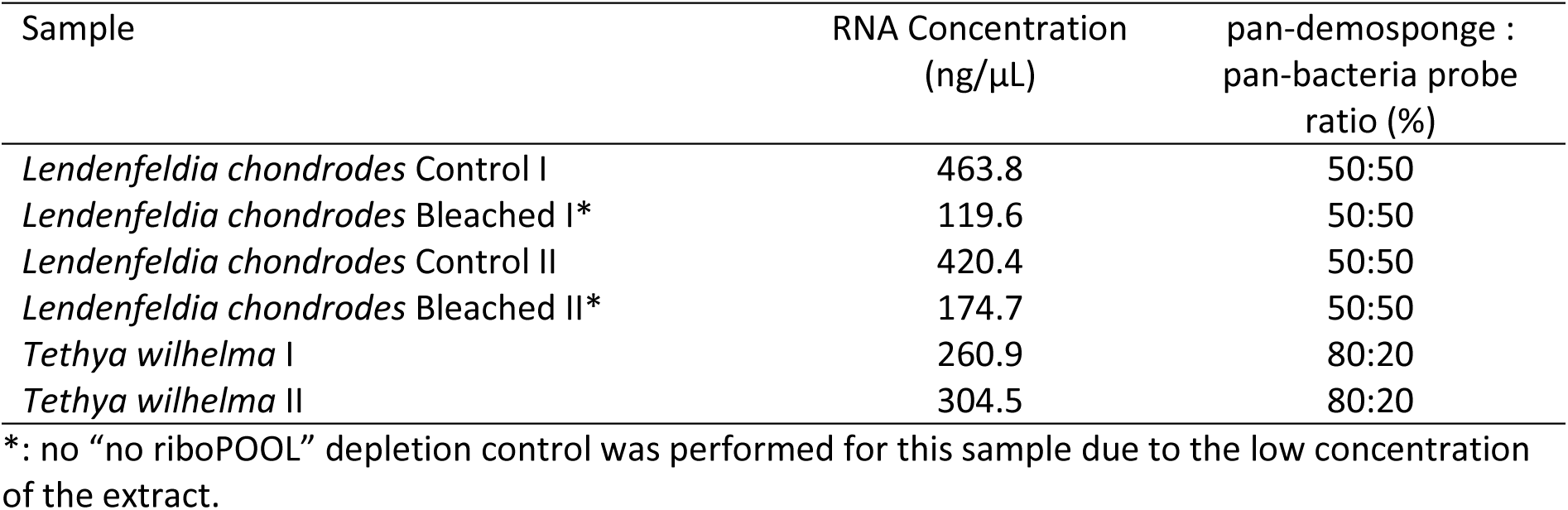
Total RNA concentration and pan-demosponge to pan-bacteria probe ratios used for rRNA depletion of *L. chondrodes* and *T. wilhelma* samples.

**Supplementary Table 3:**
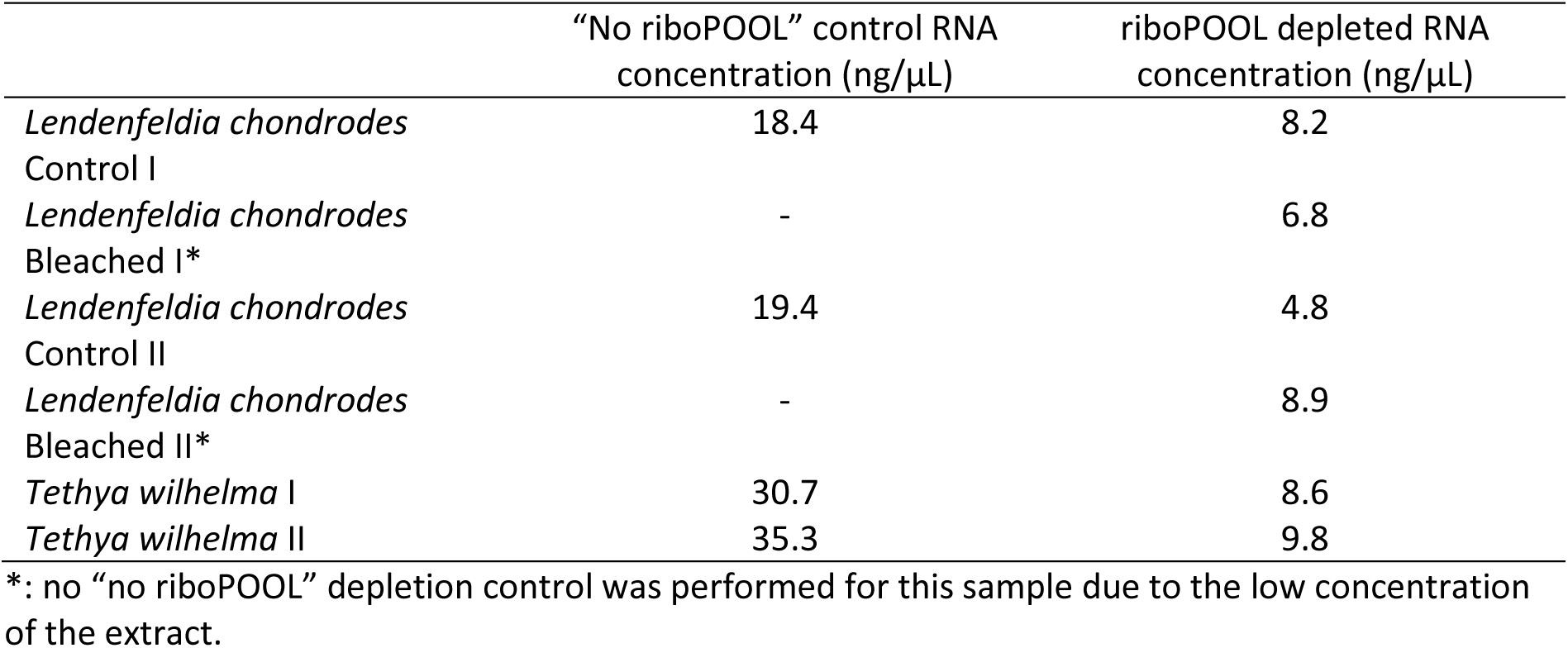
Photometrically determined concentrations of control (no riboPOOL) and depleted total RNA *L. chondrodes* and *T. wilhelma* extracts.

**Supplementary Table 4:** Cyanobacteria and Actinobacteria busco5 results for *L. chondrodes* poly-selected and rRNA-depleted transcriptomes showing the increase in detected buscos for these two phyla.

|  | Actinobacteria |  | Cyanobacteria |  |
| --- | --- | --- | --- | --- |
|  | rRNA depleted | polyA selection | rRNA depleted | polyA selection |
| Complete<br>(Single Copy,<br>Duplicated) | 42.1%<br>(15.4%, 26.7%) | 9.3%<br>(7.9%, 1.4%) | 64.1%<br>(27.9%, 36.2%) | 6.2%<br>(4.4%, 1.8%) |
| Fragmented | 5.5% | 0.7% | 6.2% | 0.4% |
| Missing | 52.4% | 90% | 29.7% | 93.4% |
| Total buscos searched | 292 |  | 773 |  |

